# Cell Type Assignments for Spatial Transcriptomics Data

**DOI:** 10.1101/2021.02.25.432887

**Authors:** Haotian Teng, Ye Yuan, Ziv Bar-Joseph

## Abstract

**Motivation:** Recent advancements in fluorescence *in situ* hybridization (FISH) techniques enable them to concurrently obtain information on the location and gene expression of single cells. A key question in the initial analysis of such spatial transcriptomics data is the assignment of cell types. To date, most studies used methods that only rely on the expression levels of the genes in each cell for such assignments. To fully utilize the data and to improve the ability to identify novel sub-types we developed a new method, FICT, which combines both expression and neighborhood information when assigning cell types.

**Results:** FICT optimizes a probabilistic function that we formalize and for which we provide learning and inference algorithms. We used FICT to analyze both simulated and several real spatial transcriptomics data. As we show, FICT can accurately identify cell types and sub-types improving on expression only methods and other methods proposed for clustering spatial transcriptomics data. Some of the spatial sub-types identified by FICT provide novel hypotheses about the new functions for excitatory and inhibitory neurons.

**Availability:** FICT is available at: https://github.com/haotianteng/FICT

## 1 Introduction

A number of different technologies have been recently developed for spatial transcriptomics. In contrast to single cell RNA-Seq most spatial transcriptomics platforms rely on image analysis by extending Fluorescence *in situ* hybridization (FISH) methods. This enables the quantification of expression levels for several genes at a single cell resolution while still recording the location of each of the cells in the sample. Examples of platforms for spatial transcriptomics include MERFISH [1,2,3], seqFISH [4,5], seqFISH+ [6], osmFISH [7], and the 3D transcriptomics record (STARmap [8]). Spatial transcriptomics techniques have now been applied to study several different organs and tissues including lung [9], kidney [10] and brain [5,6,7,8,11]. These studies have led to new insights about the set of cell types in these regions, their location and their interactions [12,13,14]

A key question in the analysis of single cell expression data (both for scRNA-Seq and for spatial transcriptomics) is the assignment of cell types. This is often the essential task performed in any analysis of such data and downstream analysis often relies on these assignments (for example, when studying cell-cell interactions [13,15]). Several packages have been developed to aid in such clustering for single cell expression data [16]. These methods often start by clustering cells (usually in low dimensional space). Next, clusters are assigned to known or new cell types based on the expression of a subset of marker genes. Most spatial transcriptomics studies have also relied on similar methods for cell type assignment. For example, in the osmFISH paper, hierarchical clustering of the gene expression profiles is used to assign cell types [7]. For the MERFISH data, cell type assignment is performed by Louvain community detection applied to a neighbourhood graph which is constructed using low dimension representation of gene expression profiles [17,18].

While using gene expression levels often leads to successful assignments, relying on scRNA-Seq cell assignment methods, for example the Seurat [19] or other clustering methods, for spatial transcriptomics may not fully utilize the available location information. Specifically, the set of neighboring cells which is known in spatial transcriptomics studies may provide valuable information about the likely cell type of a specific cell. In many cases specific cell types are known to reside together [20] or next to other types of cells [21]. Knowledge of the cell types of neighboring cells may thus provide information on the correct assignment of the cell itself. In other cases such knowledge can lead to the identification of new cell types based on their neighborhood profiles. Recently a method termed smfishHmrf was developed to utilize spatial information when assigning cell types [22]. smfishHmrf starts with an initial cell type assignment using a a support vector machine classifier which is trained using annotated expression data. Next, some assignments are updated based on a neighborhood affinity score which takes into account the fraction of cells assigned to the same cluster. While smfishHmrf utilizes some the spatial information, it only assumes that cells of the same type reside in close proximity and does not look at the overall distribution of cell types in the neighborhood of each cell. Thus, important information about the neighborhood of the cell may not be fully utilized which can lead to decrease in assignment accuracy.

To enable the use of both expression and spatial information for cell type assignment we developed FICT (FISH Iterative Cell Type assignment). FICT maximizes a joint probabilistic likelihood function that takes into account both the expression of the genes in each cell and the joint multi-variate spatial distribution of cell types. We discuss how to formulate the likelihood function and present a method for learning and inference in this model.

We applied FICT to both simulated and real spatial transcriptomics datasets. As we show using the simulation data FICT can correctly determine both expression and neighborhood parameters for different cell types improving on methods that rely only on expression levels or do not take into account the complete neighborhood of each cell. For the real data we show that the models learned by FICT for different animals for the same tissue are in good agreement, that it can indeed use the spatial information to correct errors resulting from noise in the expression values and that it can be used to identify spatially different cell sub-types even when their expression profiles are similar.

### 2 Methods

Our goal is to cluster spatial transcriptomics data using both gene expression levels and cell location. A generative mixture model is defined firstly: each cell is assigned a cell type given its neighborhood, and then the dimension reduced representation of gene expression levels are drawn from cell-type specific distribution. We next learn the parameters of this generative model by maximizing the joint likelihood of gene expression and cell location (Figure 1). The cell type is then inferred by the posterior distribution of this generative model given the gene expression level and cell location.

**Fig. 1:**
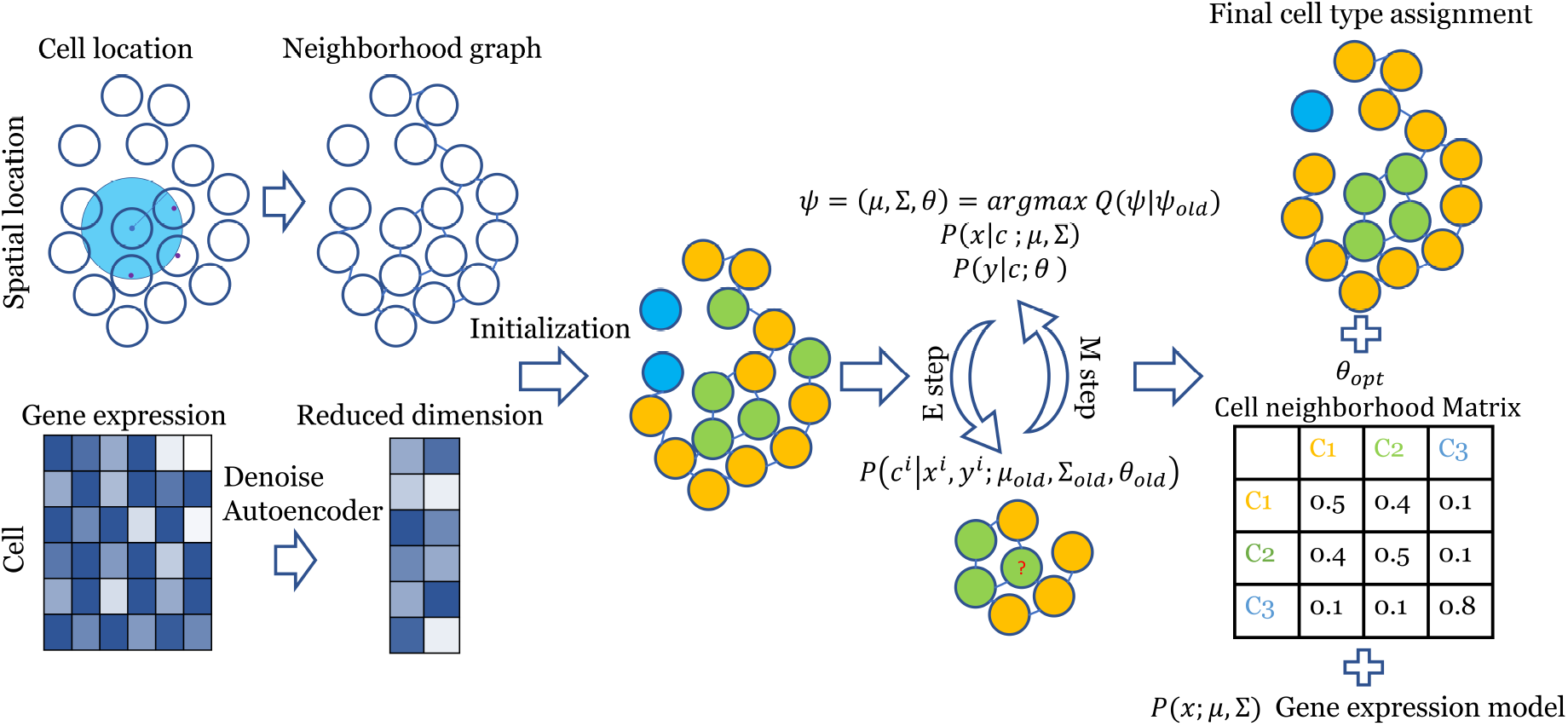
FICT pipeline. A reduced dimension expression profile is generated using a Denoising Autoencoder [23], and an undirected graph is constrcuted according to the spatial locations information. Cells are initially clustered using an expression only GMM. Next, the the model is iteratively optimized using an EM algorithm to improve the joint likelihood of the expression and neighborhood models given both the gene expression representation and the spatial graph. The final output is an assignment of cells to clusters, a Gaussian gene expression model and a Multinomial neighborhood model for each class.

### 2.1 A generative model for spatial transcriptomics data

We assume the following generative model for a single cell transcriptomics dataset *X* = *x*^(*i*)^ consisting of the gene expression levels and a graph ***G*** representing location of the cells: (1) A cell type is selected according to *P*_*θ*_(**Z**) ∝ Π_*i*_ *P* (*z*^*i*^)*ϕ*_*θ*_(*z*^*i*^, *N*_***G***_(*z*^*i*^)), in which *P* (*z*^*i*^ = *k*) = *π*_*k*_ is the prior distribution for cell type k of cell i and *ϕ*_*θ*_ is a score function with parameters *θ* capturing the relationship between neighboring cells *N*_***G***_(*c*) in ***G***. The neighbourhood cells are the cells within a threshold distance of the current cell. (2) Next, expression levels **X** are generated according to a cell type specific probability distribution *P* (*x*^(*i*)^ | *z*^(*i*)^).

Given this model the likelihood of a dataset with a set of gene expression levels *X* and cell locations (***G***) is:

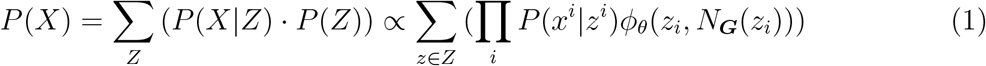

We use a multinomial distribution to model the relationship with neighborhood cells:

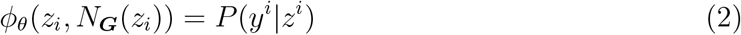

Where *y*^*i*^ is the neighborhood count of cell i, and *P* (*y*^*i*^|*z*^*i*^) follows multinomial distribution ℳ(***θ*** _*k*_). Combined, the overall likelihood function is:

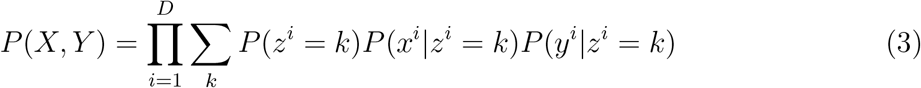

Where *X* is the dimension reduced gene expression matrix and *Y* is the neighborhood cell type count matrix for each cell, we also change the order of product and sum as y is now treated as a property of the cells. We assume that *P* (*x*^*i*^|*z*^*i*^ = *k*) follows a Gaussian distribution and *P* (*y*^*i*^|*z*^*i*^ = *k*) follows a multinomial distribution.

### 2.2 Inferring cell types (E-step)

We use an Expectation Maximization (EM) approach to learn the parameters of the model. EM iterates between the expectation (E) and maximization (M) steps. Given the generative model, to infer cell types we need to calculate the posterior probability *P* (*z*|*x, y*). However, computing these assignments is challenging since changing the assignment of a specific cell type (i.e. changes to Z’) also change the neighborhood count Y for other cells. Thus, we perform an iterative procedure as follows: In the first phase Y is treated as a fixed vector for each cell (similar to mean field approximation), and is used to calculate the posterior distribution of cell i given the gene expression matrix *x*_*i*_ and current neighborhood count *y*_*i*_ by setting:

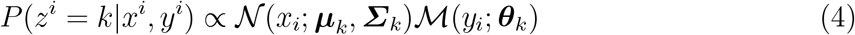

In which 𝒩(***µ***_*k*_, ***Σ***_*k*_) is a multi-variate Gaussian distribution with mean ***µ***_*k*_ and covariance matrix ***Σ*** _*k*_, and (***θ*** _*k*_) is a Multinomial distribution with ***θ*** _*k*_ as the frequency parameter, and we use *ψ* = (***µ, Σ, θ***) to denote all the model parameters. We next use the posterior distribution calculations to update cell type assignments for a subset of the cells. Specifically, we randomly select a set of non-adjacent cells in the adjacency graph ***G*** and update their types by the posterior probability. Next, the neighborhood count matrix for all cells, **Y**, is updated, and is used in the next iteration. We continue with this iterative process until convergence. This method extends the well known Iterative Condition Modes (ICM) update method [24] by updating multiple cells in each iteration instead of a single one. However, since we only update non adjacent cells, those updated cells still have the same neighborhood after each round of updates guaranteeing convergence due to the monotonical increase in overall likelihood.

### 2.3 Learning model parameters (M-step)

For M-step, we have:

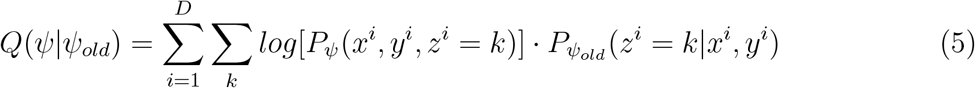

When conditioning on the cell type, the values observed for the gene expression *x*^*i*^ and neighborhood for a cell become independent. Thus, we can write:

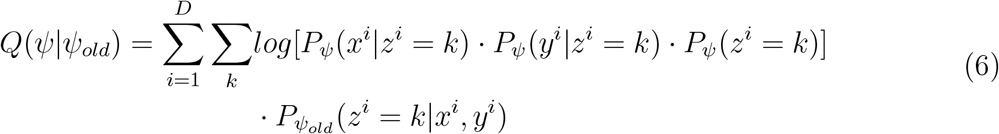

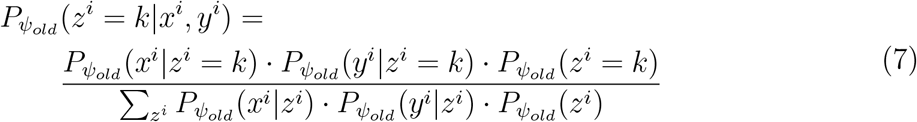

So as mentioned above (Section 2.2), the posterior distribution is calculated using an alternated ICM algorithm, in which *P* (*x*^*i*^|*z* = *k*) follows a multivariate Gaussian distribution 𝒩 (***µ***_*k*_, ***Σ***_*k*_), and the neighborhood vector for each cell *P* (*y*^*i*^|*z* = *k*) follows a Multi-Nominal distribution ℱ (***θ***)_*k*_.We set 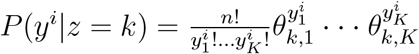, where K is the number of cell types, (*θ* _*ij*_)∈ ℝ_*K×K*_ is the neighborhood frequency of cell type j given the current cell type i, and is row-wise normalized so that | |***θ*** _*k*_ | | _1_ = 1, where ***θ*** _*k*_ is the *k*_*th*_ row of ***θ***. *π*_*k*_ = *P*_*θ*_(*z*^*i*^ = *k*) is the prior distribution for cell types.

With 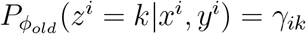 then by maximizing the given Q function, we can obtain the parameters:

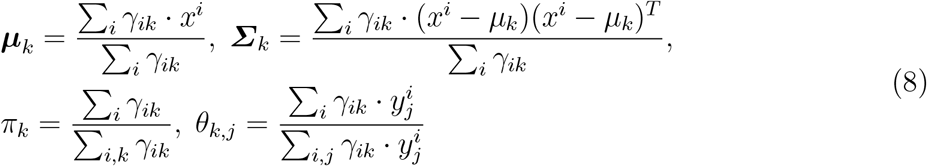

The above likelihood function assumes equal weight for each term in the two types of data (expression and neighborhood). However, there are often much more genes than cell types which can lead to over reliance on the expression data. We use two ways to address this problem, first our model is using the dimensional-reduced gene expression as input, instead of the raw expression profile. But the dimension of this input can still be high, e.g. 20, compared to the typical cell type number to be clustered, for example 7, thus then we include a weight term that balances the contribution of the gene and spatial components, named power factor (see section A1). And also during EM training, the neighborhood count is calculated in term of the assigned probability (a soft update), while usual multinomial distribution is defined in N, so we expand the scope of the multinomial distribution to ℝ to address this. See Appendix A2 for details.

### 2.4 Dimensionality reduction using denoising autoencoder

A dimension reduced representation of the original gene expression data is used as the input to our model. While the original gene expression data usually does not follow a Gaussian distribution, by using a denoising autoencoder we can transform the data to better fit such model [23]. While we use a single layer linear neural network for the auto-encoder it is possible to adapt the method to use multi-layered networks if the outcome does not fit the required Guassian distribution.

### 2.5 Generating simulated data for testing the method

We tested our method on both real and simulated data. While it is not trivial to simulate data for these experiments, simulation data provides an opportunity to test methods against ground truth which is hard to do with real data.

To obtain the simulated data we first generate a neighborhood graph, then generate a cell-type assignment on the neighbourhood graph which gives the desired neighbourhood frequency, and finally we sample expression data for each cell based on its type. See Appendix A3 for detail.

### 2.6 A comparison to Hidden Markov Random Field

Our method is a special case of a Hidden Markov Random Field (HMRF). Consider a Hidden Markov Random Field where only length 2 clique (edges) are used. For any given cell i, we have:

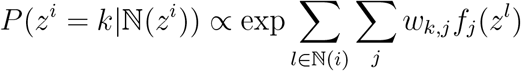

Where ℕ(*z*^*i*^) are neighbors of cell i, *W* is *K* −*by* − matrix where K is the number of cell types, and *f*_*j*_(*z*^*l*^) is a pre-defined potential function on the edges. If we set *f*_*j*_(*z*^*l*^) = **1**_*j*_(*z*^*l*^) the potential function becomes an indicator function (1 if cell *l* is of type j (*z*^*l*^ = *j*) and 0 otherwise) and so we have

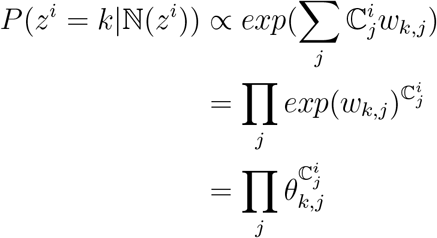

Which is the Multinomial distribution given the neighborhood count for cell i, 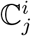. This is the same as counting the number of cells of type j that are neighbors of cell i using the frequency parameter *θ* _*k,j*_ = *exp*(*w*_*k,j*_) that we use.

## 3 Results

We developed a joint expression and location clustering method to infer cell types in spatial transcriptomics studies. To test the method we used both simulated and real single cell spatial transcriptomics data.

### 3.1 Evaluation using simulated data

While a number of spatial transcriptomics datasets exist, we do not have ground truth information about cell types in these studies. Thus, we first tested our method using simulated data where we can assign both expression and cell type and test if the method can correctly recover the cell types. As noted in Methods, generating simulated data for such analysis is not trivial since the data needs to satisfy both expression and location constraints. We developed an iterative procedure for the required spatial distributions. We used these to generate 3 datasets with different neighbourhood configurations for 3 cell types. These configurations are presented in Figure A1. We randomly sampled locations for 1,000 cells for each configuration and have also sampled expression values for each cell according to the parameters for its type.

We next tested FICT and compared it to four prior methods used to assign cell types in spatial transcriptomics data. Three of these (GMM [25,26], scanpy [27] and Seurat [19,28]) only use expression data for clustering while the fourth, smfishHmrf combines gene expression data with cell location and neighborhood information. However, unlike FICT smfishHmrf only considers neighboring cells of the same type (similar to only manually setting the diagonal values in the FICT cell neighborhood matrix and ignoring the off diagonal elements).

Results for all methods are presented in Figure 2. As can be seen, for all settings we tested FICT significantly outperforms the other methods. Seurat, which relies on shared nearest neighbor (SNN) [29] and Louvian for clustering [30] is the second most accurate method though the difference between FICT and Seurat is significant (P value ranges from 9*e*^*−*13^ to 0.028 depending on the setup).

**Fig. 2:**
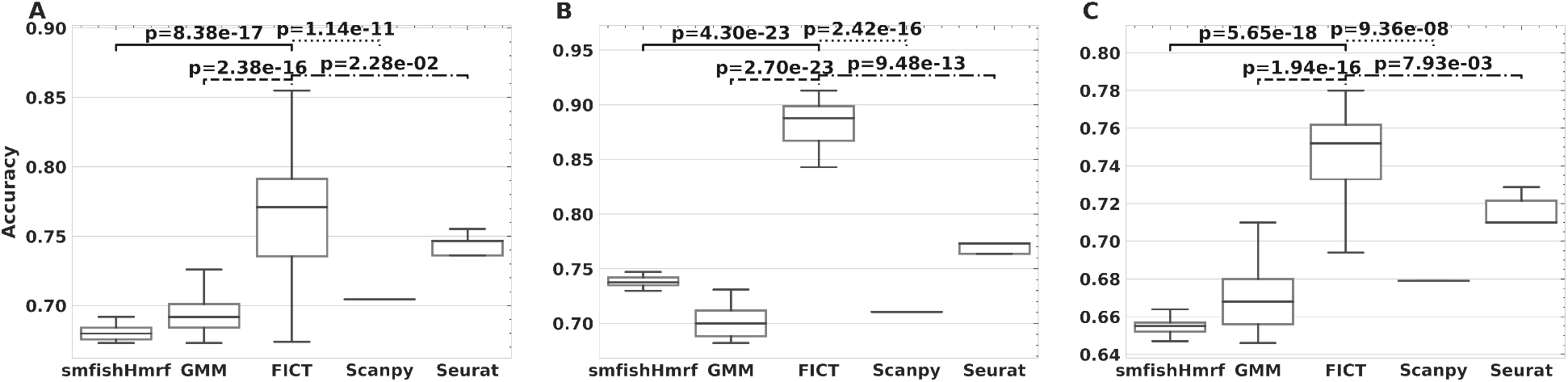
Accuracy of five different clustering methods on simulated data. We compared FICT to three expression only methods (Gaussian Mixture Models (GMM), Scanpy and Seurat) and to smfishHmrf on 3 simulation datasets generated by using different neighbourhood frequencies (see Figure A1 for distributions used for each). Accuracy was computed from 50 test runs on each dataset for each model. p value is calculated using t-test for paired samples.

### 3.2 Performance on the MERFISH dataset

We next tested FICT using real single cell spatial transcriptomics data. We first focused on mouse hypothalamus data generated by the multiplexed error-robust fluorescence *in situ* hybridization (MERFISH) method [11]. The MERFISH data profiles the expression of 258 genes in 480,000 cells from 11 animals (4 females and 7 males). Since there is no ground truth for this data, we used a different approach to compare the different clustering methods. For all gender pairs (i.e. 21 male pairs and 6 female pairs) we performed the following analysis.

Let A and B be a pair of animals from the same gender. We first train FICT on A and use the parameters learned for the model trained on A to assign cells in B. We next learn a FICT model for B. We then compare the Adjusted Rand Index (ARI) of the clustering results for the two animals. Higher ARIs mean that the results are more consistent between animals indicating better fit to the underlying biology. Note that this process is not symmetric and so results for training on A and testing on B would be different from those trained on B and tested on A.

Results for this comparison are presented in Figure 3 for both female and male animals. Note that since both Seurat and scanpy are not generative methods the models they learn on one dataset cannot be directly applied to another. Thus, for the real data we compared FICT to smfishHmrf and GMM. Results show that for 32 of the 54 pairs (59%) FICT is more consistent than GMM. The result for the larger dataset of male pairs is (29/42, 69%). The improvement upon smfishHmrf is even larger than that and FICT is more consistent in 52 of the 54 pairs (96.3%).

**Fig. 3:**
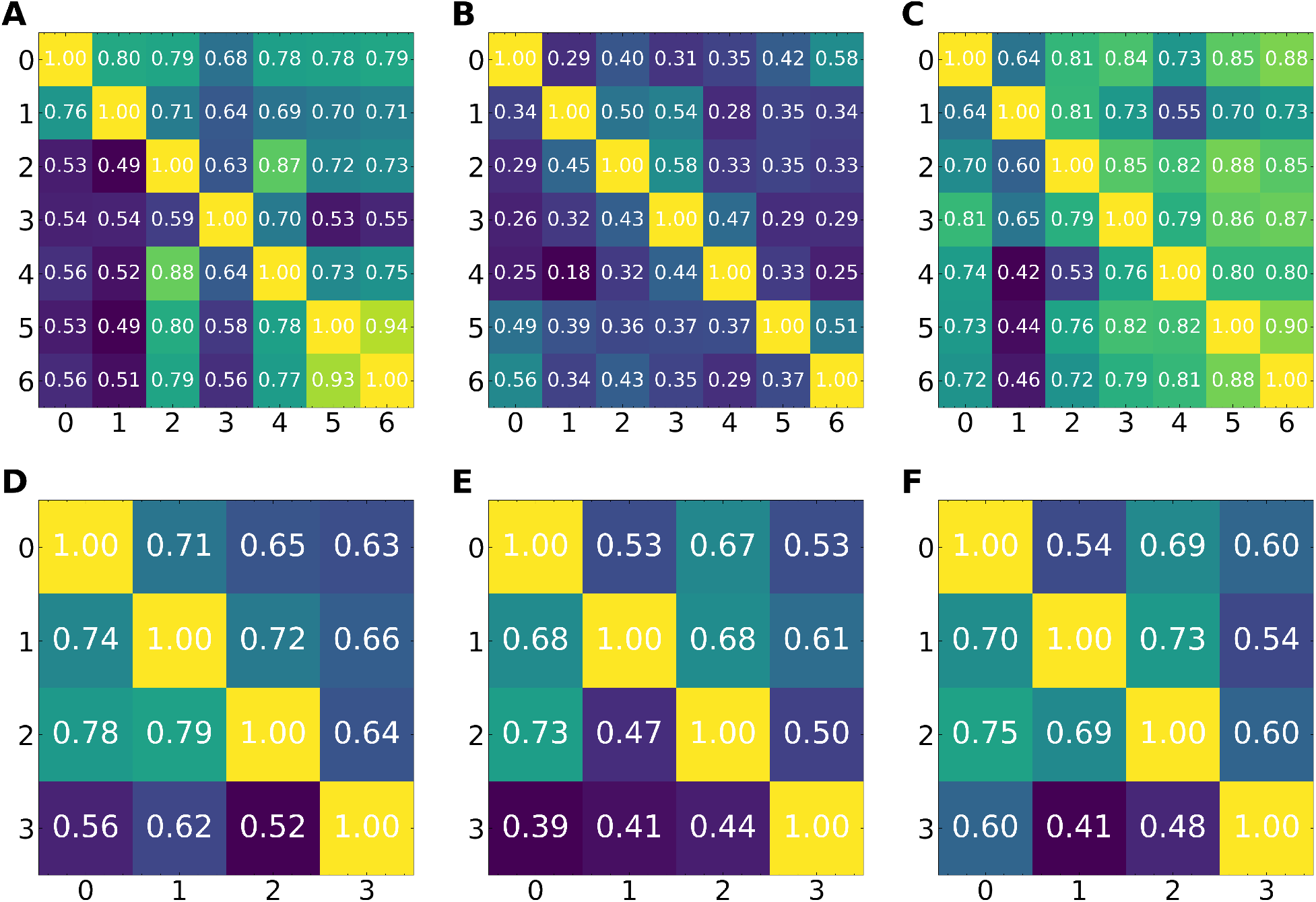
Mean Adjusted Rand index (ARI) based on cross validation analysis of the MERFISH dataset. Results presented for expression only GMM, smfishHmrf and FICT. Each entry (i,j) in the matrix represents the ARI of the two cluster assignments (one learned on animal A and applied to animal B and the other learned directly on B). (A-C) Results for the 7 Male animals (A) GMM, (B) smfishHmrf and (C) FICT. (D-F) Results for the 4 Females (D) GMM, (E) smfishHmrf and (F) FICT.

An example of the difference in assignments between expression only GMM clustering and FICT is presented in Figure 4. As can be seen, the yellow cells (Ependymal cells) are spatially clustered in the center of the hypothalamus tile profiled. However, due to small variations in gene expression, GMM assigns some cells in that cluster as OD Immature cells. In contrast FICT is able to correctly assign these cells as shown in the inset.

**Fig. 4:**
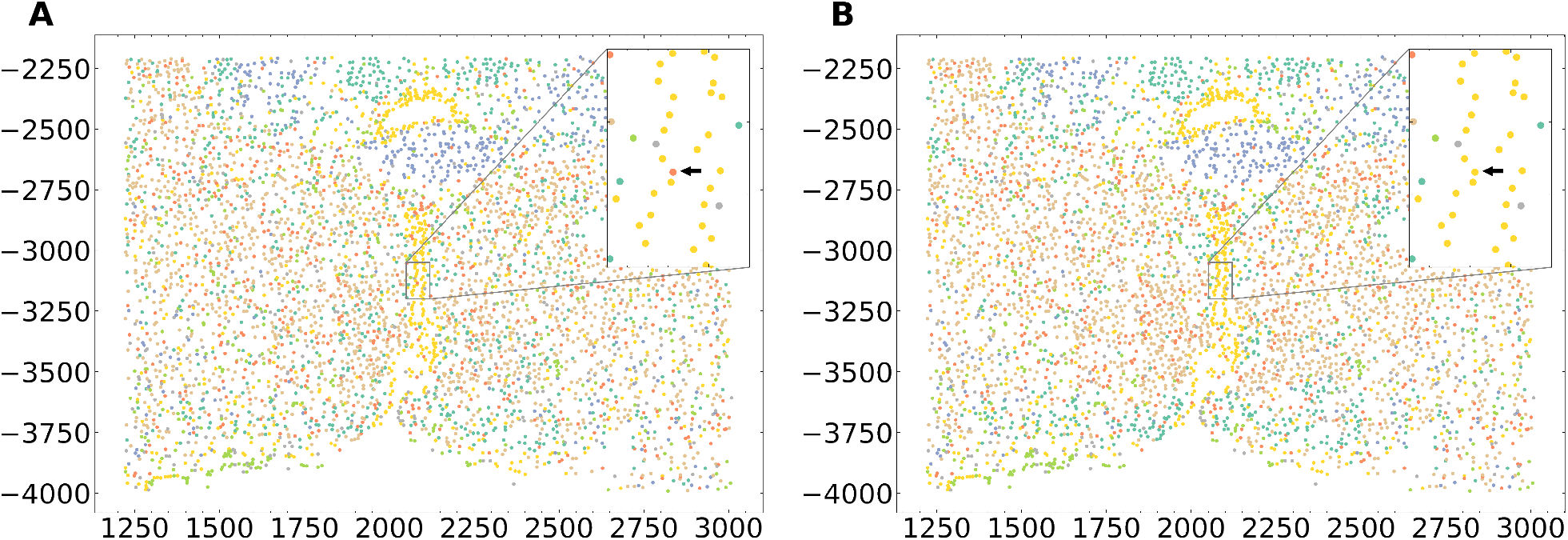
FICT can correct expression noise. Cell type assignments using expression only GMM (left) and FICT (right). Using the spatial information FICT correctly assigns Ependymal cells along the periventricular hypothalamic nucleus. In contast, the GMM method mistakenly classified the cell as OD Immature Cell.

### Sub-type clustering

An important question in the analysis of brain single cell data is the identification of new sub-types of various neuronal cells [31]. We thus examined the assignments to see if FICT can identify new subtypes of neurons. For this, we focused on the subset of excitatory neurons identified in the MERFISH dataset. FICT identified three sub-types of cells that were all determined to be excitatory in the original analysis but displayed different spatial patterns (Figure 5). To determine if the three sub-clusters are indeed different we performed differential expression (DE) analysis for each of the sub-clusters. While, as expected, their overall expression profiles are similar (leading to their similar assignment by the expression only method) we were able to identify a number of distinct genes for each of these sub-types using MAST [32]. We next performed GO enrichment analysis [33,34,35] on the significant DE genes in each sub-clusters. Results are presented in Figure 5. As can be seen, some unique functional terms are associated with each of the three sub-clusters. For example, the first sub-cluster (e0) seems to be mainly related to response to chemicals. The second (e1) seems to be related to signaling and regulation of calcium homeostasis while the third (e2) is linked to responses to activity changes and behavior. Thus, while all share similar expression profiles and act as excitatory neurons, each of the sub-clusters may have a further specific function as predicted by the spatial clustering. We performed similar sub-clustering analysis for inhibitory neurons and obtained similar results both in terms of the more coherent placing of cells from different sub-types and in terms of the unique genes and functions assigned to each of the sub-types identified by FICT (Figure 5 C and D).

**Fig. 5:**
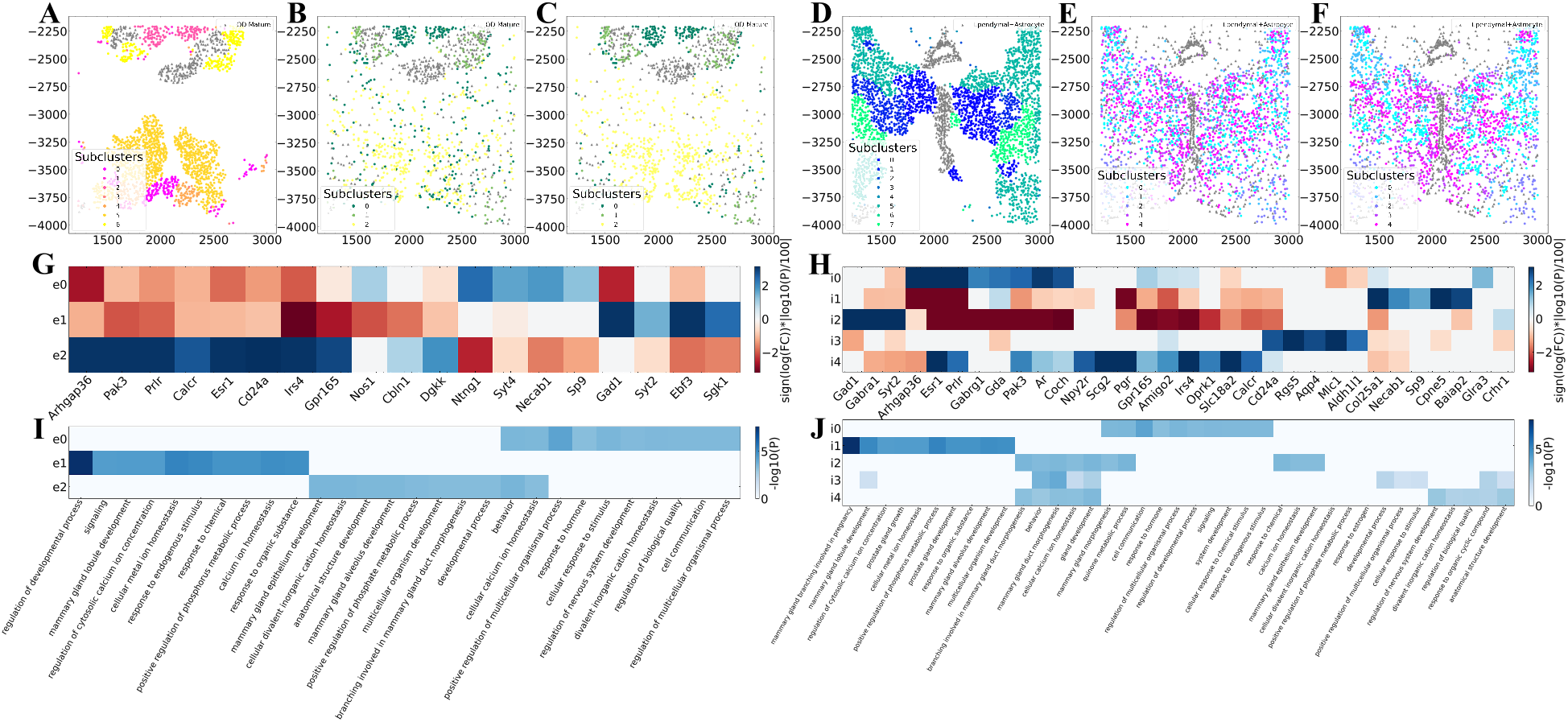
Cell subtype clustering on MERFISH data from animal 1. We used smfishHmrf (A and D), expression only GMM (B and E) and FICT (C and F) to sub-cluster excitatory neurons cells (A, B and C) and inhibitory neuron cells (D, E and F). As can be seen, for both types of neurons FICT assignments are better spatially conserved creating a central core for sub-cluster 2 surrounded by cells assigned to sub-cluster 0. In contrast, the expression only assignment mixes cells from different sub-types much more. smfishHmrf with potts model only assigns affinity score between the same cell types making it harder to infer more complex structures of synergistic activity. (E) DE genes for the three FICT sub-clusters from the excitatory neurons and (F) inhibitory neurons. As can be seen, even though the sub-clusters are overall similar in terms of their expression profiles, some genes can be identified for each of the sub-clusters. (G) **GO** enrichment analysis identifies unique functions for each of the sub-clusters on excitatory neurons and (H) inhibitory neurons.

### 3.3 Performance on osmFISH and seqFISH

To demonstrate the generality of our method we further tested it on two other datasets from two additional spatial transcriptomics platforms: osmFISH [7] and seqFISH [22]. The osmFISH dataset profiled 6,470 cells in the mouse somatosensory cortex. The seqFISH dataset profiled 1,597 cells in the mouse visual cortex. Since both datasets only profiled a single animal we performed the cross validation by manually splitting each dataset into 4 smaller regions with approximately the same number of cells. Results for these analyses are presented in Figure A4. Given the small number of cells for each dataset we see a drop in performance for all generative model methods. As the figure shows, smfishHmrf was unable to identify more than a single cell type for many of the cross validation runs resulting in errors. As for GMM and FICT while both were able to successfully assign cells in the cross validation runs for the osmFISH and seqFISH datasets, results were not as good as the MERFISH results presented above. Still, even though FICT fits more parameters than the expression only model we observe comparable performance on these smaller datasets suggesting that there is no downside to using the joint expression-spatial assignment.

## 4 Discussion

Spatial transcriptomics has emerged as a valuable tool for the analysis of single cell expression data. Similar to scRNA-Seq this technology provides information on the expression of genes at the single cell resolution. In addition, it also provides information on the location of each of the cells and their spatial relationships which can help understand cell-cell interactions, the organization of cells in specific regions and tissues and how changes in such organization impact development and disease.

A key question in spatial transcriptomics analysis is the assignment of types to the cells profiled. To date, most studies relied on the profiled expression levels for such assignment using tools and techniques originally developed for the analysis of scRNA-Seq data. While such methods work well, they do not fully utilize the information obtained in spatial transcriptomics studies. Specifically, information about the location of cells and their neighbors is usually not used in such assignments even though in several cases cell types are known to co-locate with other cells from the same or different types. To enable the use of the spatial information in cell assignments we developed FICT which uses an EM method to learn both expression and spatial distribution models. We presented a likelihood optimization function and learning and inference methods for FICT and used it to assign cell types in both simulated and real datasets.

As we have shown, for both simulated and large real datasets FICT improves on both, gene expression only methods and methods that only use part of the spatial information when assigning cell types. Since FICT estimates more parameters than expression only assignment methods its performance suffers when applied to smaller datasets. Still, even for the smallest datasets we tested on (seqFISH, which profiled only 1,500 cells) FICT performance was comparable to expression only methods making it a reasonable alternative for such methods. Since more recent studies often profile more cells, FICT is likely to generalize better to future datasets.

In addition to improved accuracy FICT can also identify cell sub-types that are similar in terms of their expression while differ in their spatial organization. As we have shown, FICT divided the set of excitatory neuron cells into three sub-types based on other cells in their neighborhood. Analysis of DE genes between these spatial clusters identified a number of biological functions that differ between the clusters indicating that each sub-type may indeed serve a different goal as predicted by FICT.

While FICT worked well for most of the datasets we tested on, there are still a number of ways in which it can be improved. We would like to improve its run-time since it currently takes one hour to perform the joint expression and spatial cell type assignment on a single animal MERFISH dataset (∼100K cells). As we noted, parts of FICT learning resemble HMRFs and so methods used to speed up HMRF inference including belief propagation can be incorporated to further improve in FICT [36]. In addition, we would like to be able to use cross validation to determine the correct number of sub-clusters to assign for each dataset rather than setting this as a user defined input.

FICT is implemented in Python and both data and an open source version of the software are available in https://github.com/haotianteng/FICT. Given the results presented in this paper we hope that it can be used to improve the analysis of the increasing number of studies that rely on spatial transcriptomics profiling.

