## Appendix for "Cell Type Assignments for Spatial Transcriptomics Data"

#### A1 Power factor

The posterior probability given by the Gaussian and Multi-Nominal distribution may be dominated by one of the two distributions and so the other information type (spatial or expression) could be neglected in the assignment updates. To overcome this we add a weight parameter to guarantee that the log posterior probability from both distributions will have equal mean. This leads to the following changes to the likelihood function:

$$G^{\alpha}(x, \mu, \Sigma) = \frac{exp(-\frac{1}{2}(x-\mu)^T \alpha \Sigma^{-1}(x-\mu))}{\sqrt{(2\pi)^k (1/\alpha)^k |\Sigma|}}$$

$$\mathcal{M}^{\boldsymbol{\beta}}(y, \theta) = \frac{\Gamma(\sum_{i=1}^{k} y_i \beta_i + 1)}{\prod_{i=1}^{k} \Gamma(y_i \beta_i + 1)} \prod \theta_i^{y_i \beta_i}$$

Here $\alpha$ is the factor for gene model and $\beta$ is the factor for spatio model, $\Gamma(x)$ is the Gamma function. Using the weight the MLE of the covariance $\boldsymbol{\Sigma}$ and frequency parameter $\theta$ becomes:

$$\boldsymbol{\Sigma}_k = \frac{\sum_i \gamma_{ik} \cdot \alpha(x^i - \mu_k)(x^i - \mu_k)^T}{\sum_i \gamma_{ik}}, \ \theta_{k,j} = \frac{\sum_i \gamma_{ik} \cdot y_j^i \beta_j}{\sum_{i,j} \gamma_{ik} \cdot y_j^i \beta_j} \tag{9}$$

The MLE of mean $\mu$ stays the same.

#### A2 Analytical continuation of Multinomial distribution

During EM training and in the power factor model, we use the following form for the multinomial distribution:

$$\mathcal{M}(y, \theta) = \frac{\Gamma(\sum_{i=1}^{k} y_i + 1)}{\prod_{i=1}^{k} \Gamma(y_i + 1)} \prod \theta_i^{y_i} \tag{10}$$

The correctness of the expression can be shown if $y_i$ is an integer, and since $\Gamma(y_i + 1) = y_i!$

$$\mathcal{M}(y, \theta) = \frac{(\sum_{i=1}^{k} y_i)!}{\prod_{i=1}^{k} y_i!} \prod \theta_i^{y_i} \tag{11}$$

Which is the original definition of multinomial distribution.

#### A3 Cell type assignment in simulation

It is not trivial to generate a cell-type configuration whose neighbourhood frequency follows a given multinomial distribution. We have used the following strategy to address this problem: First we generate the 2-D physical location matrix for each cell $\mathbf{Q} \in \mathbb{R}_{N\times 2}$. We use this and a distance threshold $D$ to generate the neighborhood graph $\boldsymbol{G}$. Then we assign an initial cell type to each cell by sampling from the prior distribution $\mathbf{P_c}$. Next, we apply two sampling methods, i.e., Gibbs sampling and annealing simulation, to approximate the cell type assignment. However before we assigning the cell type, we notice here that not all target neighborhood probability matrix $\mathbf{A}$ are valid, for example the following given matrix is not a valid neighborhood frequency matrix:

$$\begin{bmatrix} 0.2\,0.8\,0.0 \\ 0.1\,0.3\,0.6 \\ 0.2\,0.5\,0.3 \end{bmatrix}$$

Here row i of the matrix $\mathbf{A}_i$ represent the probability of the cell types of the neighborhood for a given cell type i, i.e. $\mathbf{A}_{i,j}$ represent the probability of cell type j being the neighbourhood of cell type i. The above given matrix is invalid because the probability of cell type 3 being neighborhood of cell type 1 can not be 0 ($\mathbf{A}_{1,3}$) as the cell type 1 being neighborhood of cell type 3 ($\mathbf{A}_{1,3}$) is not zero.

So we have to determine if a given probability matrix is a valid neighborhood matrix, an observation is that count matrices should be symmetric, since if cell i is a neighbour of cell j, then the reverse is also true, any valid probability matrix is the normalization of a count matrix. Based on this observation we use the following criteria to find a close approximating neighbourhood matrix if the given probability matrix is not valid: if we can find a diagonal matrix $\mathbf{D} = \{d_{11}, ..., d_{nn}\}$, so that $\mathbf{D} \cdot \mathbf{A}$ is a symmetric matrix, then the matrix $\mathbf{A}$ is valid. We use the following algorithm to find an approximating valid neighbourhood matrix $\mathbf{A}^*$ from a given probability matrix $\mathbf{A}$.

**Algorithm 1:** Finding valid neighborhood probability matrix

**Result:** A matrix $\mathbf{A}^*$ that is the closest matrix of the given probability matrix $\mathbf{A}$
**Initial:** A initial diagonal matrix $\mathbf{D}$
$\mathbf{D}_{op} = \arg\min_{\mathbf{D}} \|\mathbf{D} \cdot \mathbf{A} - \mathbf{A}^T \cdot \mathbf{D}\|_2 + \lambda \cdot (\|\mathbf{D}\|_1 - 1)^2$
$\mathbf{C}^* = (\mathbf{D}_{op} \cdot \mathbf{A} + \mathbf{A}^T \cdot \mathbf{D}_{op})/2$
$\mathbf{A}^* = Normalize(\mathbf{C}^*)$

After obtaining the approximated target neighborhood frequency matrix $\mathbf{A}^*$, we first apply Gibbs sampling, update each cell by sampling from the posterior probability given its neighborhood and the target neighborhood frequency $\mathbf{A}*$. In each update, a cell type neighborhood probability matrix $\mathbf{A}^{(t)}$ is computed. During annealing simulation, in each step we attempted to swap the cell type of two randomly selected cells, and the KL-divergence between the current neighborhood frequency $\mathbf{A}^{(t)}$ and the target neighborhood frequency $\mathbf{A}*$ is recorded, the swap is accepted if the resulted KL-divergence is smaller otherwise rejected by a chance proportional to the increase of KL-divergence. We firstly apply Gibbs sampling until converge and then apply annealing sampling until converge.

Next, based on the cell type assigned by the above procedure, we generate gene expression values for each cell based on its assigned type. Each cell type has approximately the same number of cells. 30% of genes for each type are cell-type specific, the rest follow a global pattern, to mimic, for example, cell cycle and other cellular activity related genes. Gene expression is assumed to follow a zero-inflated log-normal distribution. While some studies use zero-inflated negative binomial distribution for scRNA-Seq count data simulations[37], spatial data is image based and so continuous. As we mentioned before, a more sophisticated dimensional-reduced technique can be used to further extend the power of the gene model, and may give better accuracy, but it's out of the scope of this paper.

**Algorithm 2:** Assigning cell-types using Gibbs Sampling

**Result:** A cell type count matrix $\mathbf{C}^*$ and a cell type assignment vector $\mathbf{T}^*$
**Initial:** A initial cell type assignment $\mathbf{T}^{(0)}$, time step t=0
**Input :** A target probability matrix $\mathbf{A}^*$
**while** $D_{KL}(\mathbf{A}^{(t)} \| \mathbf{A}^*) > desireddistance$ **do**
  $\mathbf{C}^{(t)} = 0$;
  **for** *i cell* $c_i \subset$ *all cells* **do**
    $\mathbf{T}^{(t)}[i] = \text{Assign}(\mathbb{N}(c_i), \mathbf{A}^*, P_{prior}(c_i))$
  **end**
  **for** *i cell* $c_i \subset$ *all cells* **do**
    $t_i = \mathbf{T}^{(t)}[i]$;
    $\mathbf{C}^{(t)}[t_i, :] \mathrel{+}= \text{Count}(\mathbb{N}(c_i))$;
  **end**
  $\mathbf{A}^{(t)} = Normalize(\mathbf{C}(t))$;
**end**
**Function** `Assign`$(\mathbb{N}(c_i), \mathbf{A}, P_{prior}(c_i)^*)$**:**
  #Input Args;
  #$\mathbb{N}(\mathbf{x})$: A vector indicate the cells that is directly connected to cell in $\mathbf{x}$;
  #$P_{prior}$: A vector indicate the prior distribution of given cell type;
  #$\mathbf{A}^*$: The target neighborhood probability matrix.;
  $P(c_i|c_{-i}) = P(c_i|\mathbb{N}(\mathbb{N}(c_i))) = \frac{P(\mathbb{N}(c_i)|c_i)^{1/s} \cdot \prod_{c_j \in \mathbb{N}(c_i)} P(\mathbb{N}(c_j)|c_j)^{1/s}}{\sum_{c_i} P(\mathbb{N}(c_i)|c_i)^{1/s} \cdot \prod_{c_j \in \mathbb{N}(c_i)} P(\mathbb{N}(c_j)|c_j)^{1/s}}$;
  # s is a soft factor ($s > 1$);
  $P(\mathbb{N}(x)|x) = (\sum \text{Count}(\mathbb{N}(x)))! \prod_j \frac{\mathbf{A}[x][j]^{\text{Count}(\mathbb{N}(x))_j}}{\text{Count}(\mathbb{N}(x))!}$;
  Assign $c_i$ according to $P(c_i|c_{-i})$;
  return $c_i$

#### A4 Comparison to HMRF

As mentioned in the main text our method can be thought of as a variant method of the Hidden Markov Random Field (HMRF). Consider a Hidden Markov Random Field where only length 2 clique (edge) is take into account, for any given cell i, we have:

$$P(z^i = k|\mathbb{N}(z^i)) \propto \exp \sum_{l \in \mathbb{N}(i)} \sum_j w_{k,j} f_j(z^l)$$

Where $\mathbb{N}(z^i)$ are neighborhood cells of cell i, $W$ is $K - by - K$ matrix where K is the number of cell type, $f_j(z^l)$ is some arbitrary function. Commonly, if we select $f_j(z^l) = \mathbf{1}_j(z^l)$ which is an indicator function which equals to 1 if cell $l$ is as type j ($z^l = j$) otherwise 0, we will have

$$P(z^i = k|\mathbb{N}(z^i)) \propto exp(\sum_j \mathbb{C}^i_j w_{k,j}) = \prod_j exp(w_{k,j})^{\mathbb{C}^i_j} = \prod_j \theta_{k,j}^{\mathbb{C}^i_j}$$

Which gives the Multinomial distribution given the neighborhood count of cell i, $\mathbb{C}^i_j$, the number of cells in type j that is a neighborhood of cell i with the frequency parameter $\theta_{k,j} = exp(w_{k,j})$

### A5 Supplementary Tables and Figures

| | Addictive | Exclusive | Consecutive |
|---|---|---|---|
| GMM | 0.695± 0.005 | 0.701 ± 0.021 | 0.671 ± 0.019 |
| smfishHmrf | 0.681± 0.006 | 0.734 ± 0.040 | 0.655 ± 0.005 |
| FICT | **0.764** ± 0.039 | **0.865** ± 0.069 | **0.738** ± 0.035 |
| Scanpy | 0.701 ± 0.023 | 0.672 ± 0.005 | 0.634 ± 0.100 |
| Seurat | 0.743 ± 0.006 | 0.770 ± 0.004 | 0.715 ± 0.008 |

Table A1: Accuracy of five cell type assignment methods on 3 simulation datasets generated from different neighbourhood frequency (Figure A1). FICT performs significantly better on all configurations. Accuracy is calculated as the highest hit percentage obtained by permuting the assigned cluster label and compare it to the true label.

##### A5.1 Neighbourhood Frequency of three simulation datasets

The three different negihbourhood frequency configurations in simulation are shown in Figure A1.

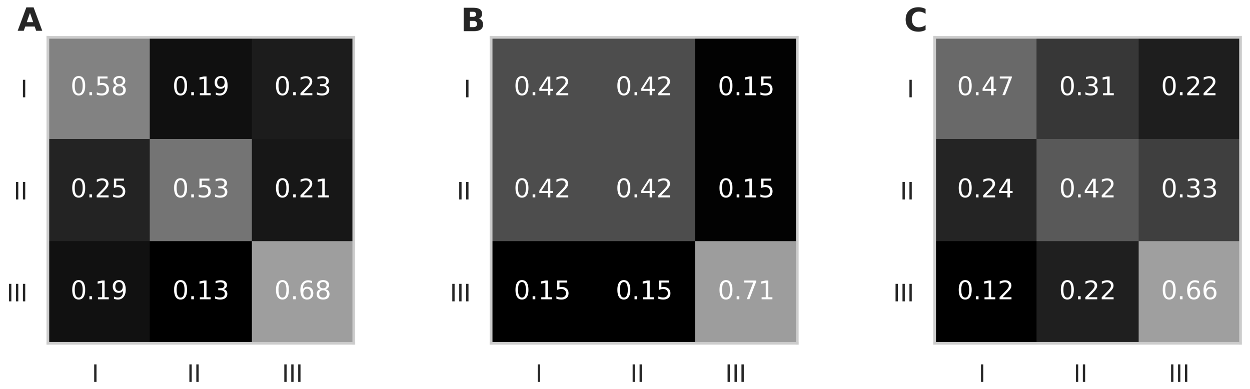


Fig. A1: The neighbourhood matrix in the corresponding simulation dataset in Figure2, the $i_{th}$ row represent the neighbourhood frequency of cell type i (i=1,2,3). (A) An addictive configuration where each cell type tend to appear around its own kind. (B )An exclusive configuration, in which type 1 cell and type 2 cell are more likely to appear beside each other, wheres type 3 cell is exclusive. (C) A consecutive configuration, where type 2 cell is likely to appear around type 1 cell, type 3 cell is likely to appear around type 2 cell, while type 3 cell is unlikely to appear around type 1 cell.

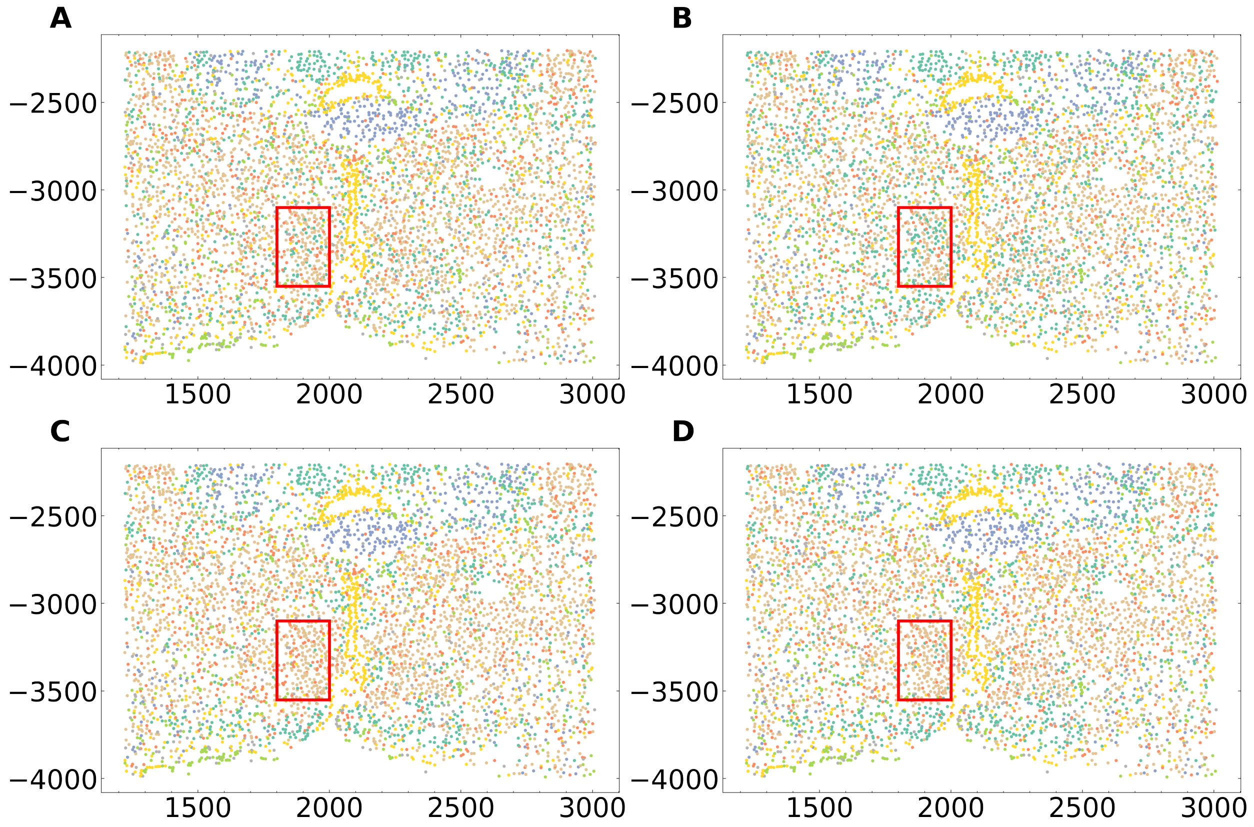


Fig. A2: Illustrations of the clustering result on MERFISH dataset female animal 1 by (A) GMM trained on animal 1. (B) GMM trained on animal 2. (C) FICT trained on animal 1. (D) FICT trained on animal 2. The Ependymal cell colored by yellow form a ladle-like pattern, which is known to be the periventricular hypothalamic nucleus, and is clearly addressed. Red bounding box address a region where cell type assignments are different between the 4 models, compare to GMM, FICT gives more consistent result in this region.

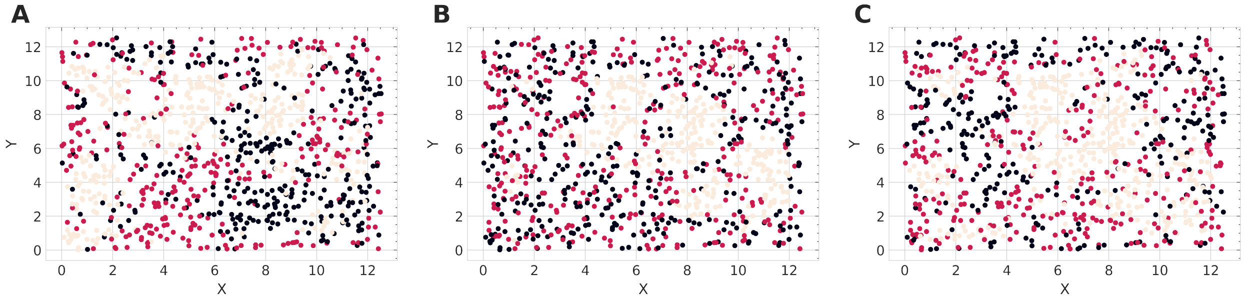


Fig. A3: A scatter plot showing the cell distribution of three different simulation configuration. (A) Addicitive configuration where cells prefer to aggregate with cells from same type. (B) Exclusive configuration where type 1 and type 2 cells (black and red cells) prefer to aggregate together but keep away from type 3 cell (yellow cells) (C) Consecutive configuration, type 1 cell (black cells) tends to surround type 2 cells (red cells) but not type 3 cells (yellow cells), type 2 cells (red cells) aggregate with both type 1 and type 2 cells (black and red cells) and type 3 cells (yellow cells) tends to surround type 2 cells (red cells) but not type 1 cells (black cells), this configuration causing the formation of stripe pattern.

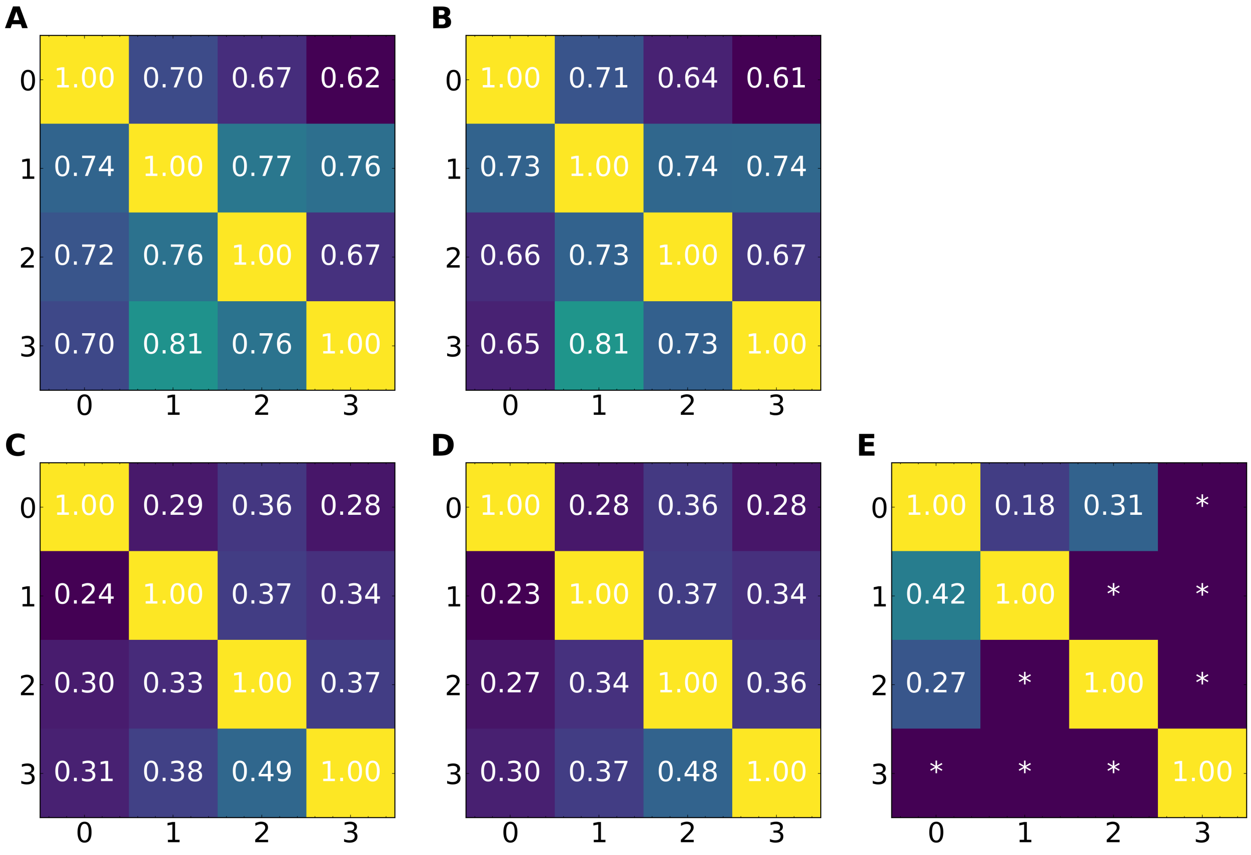


Fig. A4: Cross validation score of (A) GMM and (B) FICT on osmFISH, (C) GMM, (D) FICT and (E) smfishHmrf on seqFISH. * Denote that this cross validation using the corresponding sub-datasets failed because smfishHmrf has only identified a single cell type in the cross validation dataset resulting in an error message. Similarly, cross validation using smfishHmrf failed for all osmFISH comparisons.

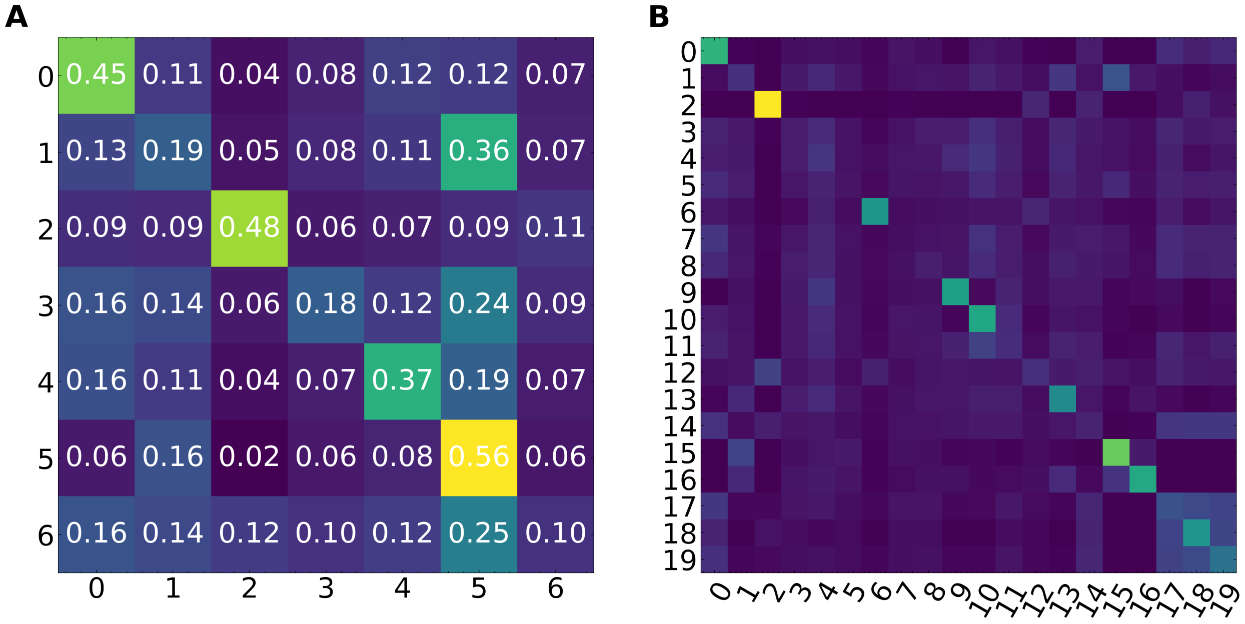


Fig. A5: Heatmap of the multinomial model parameter demonstrate the neighbourhood frequency of different cell types. (A) is from 7-Class model and (B) is from 20-Class model trained using MERFISH dataset.
